# Neurovascular Impulse Response Function (IRF) during spontaneous activity differentially reflects intrinsic neuromodulation across cortical regions

**DOI:** 10.1101/2024.09.14.612514

**Authors:** Bradley C. Rauscher, Natalie Fomin-Thunemann, Sreekanth Kura, Patrick R. Doran, Pablo D. Perez, Kıvılcım Kılıç, Emily A. Martin, Dora Balog, Nathan X. Chai, Francesca A. Froio, Patrick F. Bloniasz, Kate E. Herrema, Rockwell Tang, Scott G. Knudstrup, Andrew Garcia, John X. Jiang, Jeffrey P. Gavornik, David Kleinfeld, Michael E. Hasselmo, Laura D. Lewis, Sava Sakadzic, Lei Tian, Gal Mishne, Emily P. Stephen, Martin Thunemann, David A. Boas, Anna Devor

## Abstract

Ascending neuromodulatory projections from deep brain nuclei generate internal brain states that differentially engage specific neuronal cell types. Because neurovascular coupling is cell-type specific and neuromodulatory transmitters have vasoactive properties, we hypothesized that the impulse response function (IRF) linking spontaneous neuronal activity with hemodynamics would depend on neuromodulation.

To test this hypothesis, we used optical imaging to measure (1) release of neuromodulatory transmitters norepinephrine (NE) or acetylcholine (ACh), (2) Ca^2+^ activity of local cortical neurons, and (3) changes in hemoglobin concentration and oxygenation across the dorsal surface of cerebral cortex during spontaneous neuronal activity in awake mice.

A canonical convolution model with a global, stationary IRF (i.e., the convolution kernel) describing evolution of total hemoglobin (HbT, reflective of dilation dynamics) with respect to Ca^2+^, resulted in a poor fit to the data. However, the HbT time-course was well predicted, pixel-by-pixel, by a weighted sum of Ca^2+^ and NE time-courses. Consistent with this result, modeling HbT as a weighted sum of stationary Ca^2+^ - and NE-specific IRFs (IRF_Ca2+_ and IRF_NE_) convolved with the respective time-courses dramatically improved the fit compared to the global IRF. IRF_Ca2+_ and IRF_NE_, estimated from the data, were positive and negative, respectively. In contrast to NE, ACh was largely redundant with Ca^2+^ and therefore did not improve HbT estimation. Because NE covaried with arousal, we observed instances of the diminished hemodynamic coherence between cortical regions during high arousal despite coherent behavior of the underlying neuronal Ca^2+^ activity.

We conclude that while neurovascular coupling with respect to neuronal Ca^2+^ is a dynamic and seemingly complex phenomenon, hemodynamic fluctuations can be captured by a simple linear model with stationary IRFs with respect to the underlying dilatory and constrictive forces. In the current study, these forces were captured by the positive IRF_Ca2+_ (dilation) and negative IRF_NE_ (constriction). Without accounting for NE neuromodulation and the associated vasoconstriction, diminished hemodynamic coherence, commonly referred to as “functional (dys)connectivity” in BOLD fMRI studies, can be falsely interpreted as neuronal desynchronizations.

## Introduction

Ascending adrenergic and cholinergic projections from the locus coeruleus (LC) and basal forebrain (BF) have long been associated with shaping information processing in local and large-scale cortical networks (1, 2) and modulation of cognitive functions such as arousal and attention (3). The respective neurotransmitters, norepinephrine (NE) and acetylcholine (ACh), are vasoactive affecting cerebral blood flow (CBF) in two ways: directly, by acting on smooth muscle of arterioles, and indirectly, via modulation of local neurons (4, 5). These neuromodulatory systems are active in any conscious human subject participating in a Blood Oxygenation Level Dependent (BOLD) functional Magnetic Resonance Imaging (fMRI) study. And yet, how these systems affect the relationship between activity of local neuronal circuits and hemodynamics across cortical regions remains unclear.

Optogenetic (OG), chemogenetic, and pharmacological perturbations of NE and ACh signaling in animals produce wide-spread hemodynamic changes and modulate the temporal correlation of hemodynamics across brain regions (i.e., functional connectivity, FC) (6–13). Complementary to the perturbation approach, it is now possible to directly visualize release of neuromodulatory transmitters in cerebral cortex *in vivo*, thanks to the recent development of genetically encoded optical biosensors (14–17) and wide-field, large-scale “mesoscopic” optical imaging (18–20).

In the present study, we leveraged our version of a wide-field mesoscope (18), as well as 2-photon microscopy, to assess the relationship between Ca^2+^ activity of cortical neurons and the associated hemodynamics during spontaneous activity in fully awake mice. We show that while this relationship varied across cortical regions and NE levels, hemodynamic fluctuations were accurately predicted given the local Ca^2+^ and NE signals. During high arousal, the dynamic nature of IRF resulted in diminished hemodynamic coherence between certain cortical regions despite coherent behavior of the underlying neuronal Ca^2+^ activity.

## Methods

### Experimental animals

All experimental procedures were performed in accordance with the guidelines established by the Boston University Institutional Animal Care and Use Committee. Standard rodent chow and water were provided ad libitum. Mice were housed in a 12-hour light cycle with lights turning on at 7:30 AM. We used 15 adult mice of either sex from the Thy1-jRGECO1a GP8.20 line (21). Cortex-wide expression of GRAB_NE_ or GRAB_ACh3.0_ was induced by either (a) injection of 1.5 µL AAV9-hSyn-GRAB_ACh3.0_ (or AAV9-hSyn-GRAB_NE2m_) (WZ Biosciences or BrainVTA, 3.06×10^13^ GC/mL) into each transverse sinus (3 µL per animal) in 1-day old neonates (20) or (b) retro-orbital virus injection of PHP.eB variants (BrainVTA, PT-1344, rAAV-hSyn-NE3.1,AAV2/PHP.eB; 2×10^13^ vg/mL, 50 µl used) in adult (6 to 9-week old) mice (22).

### Animal procedures

The surgical procedure was modified from that previously described (23, 24). Dexamethasone was injected ∼4 h prior to surgery (IP 4.8 mg/kg at 4 mg/ml concentration) to prevent brain swelling due to surgical intervention. In addition, slow-release Buprenorphine and Meloxicam were injected before surgery. During surgery, mice were anesthetized with either isoflurane (2% in O_2_ initially, 1% in O_2_ during all procedures), ketamine/xylazine or a mixture of medetomidine, midazolam and fentanyl (MMF), and their body temperature was maintained at 37°C. A custom-designed headpost machined from titanium was attached to the cranium. Two pieces of curved or flat glass were used to replace the dorsal cranium, one per hemisphere (24). To prevent heat loss through the window, either silicone or cotton was placed on top of the glass window and contained with a protective 3D-printed cap fixed to the headpost. Mice were allowed to recover for at least one week following surgery before habituation to head fixation and at least 3 weeks before experiments.

### Behavioral training and recording

Starting at least 1 week after the surgical procedure, mice were habituated in 1 session/day to accept increasingly longer periods of head restraint (up to 2 hrs). During the head restraint, the animal was placed on a suspended bed. A drop of sweetened condensed milk was offered a few times during the fixation as a reward. Habituated head-fixed mice consumed the reward milk and often exhibited periods of putative sleep defined as the eye pupil constriction accompanied by a large-amplitude, prolonged vasodilation (25). Mice were free to readjust their body position and from time to time displayed natural grooming behavior. In both mesoscopic and 2-photon experiments, a CCD camera (Basler, acA1920-150uc) with a variable zoom lens (Edmund Optics, 67715) was used for continuous observation of the mouse face during imaging. In 2-photon experiments, we used a 940 nm LED (Thorlabs, M940L3) to illuminate the face of the animal, and a 920-nm long pass filter to prevent illumination light from reaching PMT detectors. In 1-photon experiments, we used a 940 nm LED (Thorlabs, M940L3) to illuminate the face of the animal. The camera frames were synchronized with image acquisition and recorded. In addition, an accelerometer (Analog Devices, ADXL335) was placed underneath the mouse bed to record animal’s motion. Periods of putative sleep were excluded from data analysis.

### Mesoscopic imaging

Widefield imaging was performed using a home built mesoscope described in (18) (**Fig. 1A**). A single sCMOS camera coupled to an objective lens (Olympus ZDM-1-MVX063) was used to sequentially record four channels: a green fluorescence channel for imaging of GRAB_NE_ or GRAB_ACh_, a red fluorescence channel for imaging of jRGECO1a, and two reflectance channels at 525 nm and 625 nm for quantification of oxy-, deoxy-, and total hemoglobin (HbO/HbR/HbT). A 10x10 mm^2^ field of view covered the entire dorsal cortex. Images were acquired at 10 Hz, with a spatial resolution of 20 μm at 4x4 binning. Fluorescence excitation was achieved through epi-illumination. For GRAB excitation, a 470-nm LED (ThorLabs SOLIS-470C) was used with a 466/40 nm bandpass filter (Semrock). For jRGECO1a excitation, a 565-nm LED (ThorLabs SOLIS-565C) was used with a 560/14-nm bandpass filter (Semrock) and an additional 450-nm longpass filter (Chroma) to eliminate a secondary emission peak around 425 nm. The emission pathway included a dual-band dichroic mirror (488/561-nm, Semrock) and a multi-band filter (523/610-nm, Semrock). This configuration allowed GRAB and jRGECO1a fluorescence (500-540 nm and 580-640 nm, respectively) to reach the detector while blocking excitation light (440-480 nm and 550-570 nm). For imaging of reflectance, illumination LEDs were positioned in a ring around the objective, and an aluminum hemisphere was placed between the objective and cranial window to provide diffuse illumination (26). The estimated changes in HbO and HbR were used to correct fluorescence signals for the hemodynamic artifact (27).

**Figure 1:**
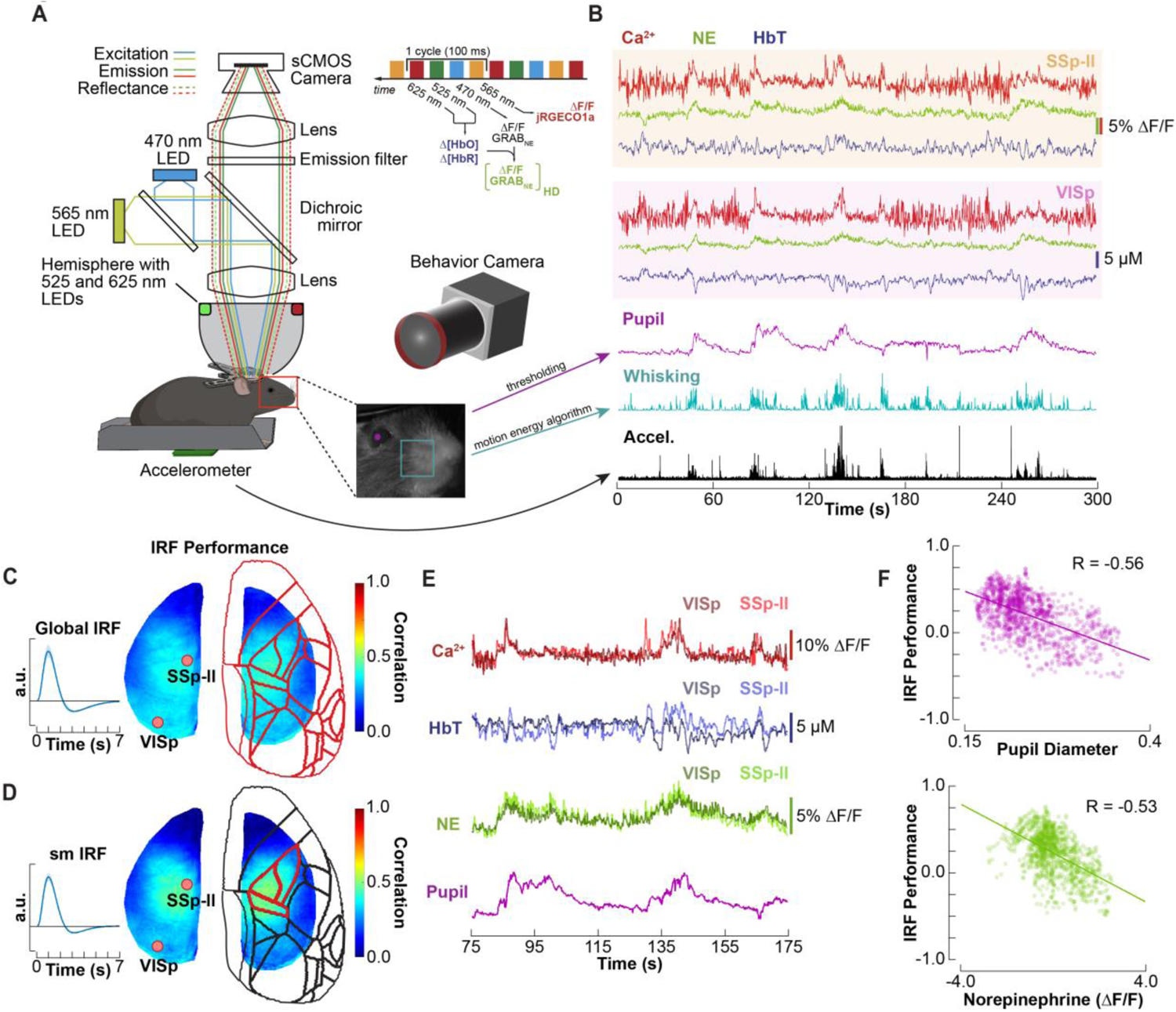
IRF linking spontaneous hemodynamics to neuronal Ca^2+^ varies across cortical space and time. **A)** Schematics of the mesoscopic (wide field) imaging setup. Upper right: timing of frame acquisition by the CMOS camera. Lower right: an image of mouse face acquired by the CCD camera for quantification of behavior: eye pupil, whisking, and body movement. **B)** Example Ca^2+^, NE, and HbT time-courses derived from two cortical regions, SSp-ll and VISp (32) along with behavior readouts. **C)** Global IRF estimation from Ca^2+^ and hemodynamic data and the IRF performance quantified as the correlation between the estimated and experimental HbT (n=15 mice). **D)** Same as (C) but using only the somatosensory cortex (SSp-tr, SSp-ll, and SSp-ul; (32)) for estimation of the IRF (n=15 mice). **E)** Example Ca^2+^, NE, and HbT time-courses from SSp-ll and VISp. **F)** IRF Accuracy of HbT prediction with the IRF from (D) including the data shown in (B) quantified as the correlation between the estimated and experimental HbT (n=1 mouse, 3 runs). Each point on the scatter plot corresponds to a correlation coefficient for a 15-s sliding window. The correlation coefficient was calculated for each pixel, averaged within the SSp-bfd region, and then plotted against the average pupil diameter (top) or NE (bottom) within the same time window.

Data were acquired in 10-min “runs.” Multiple runs were acquired on each experimental day for one mouse, and each mouse was imaged on multiple days (**Suppl. Table 1**).

### Two-photon imaging

The GRAB sensor (GRAB_NE_ or GRAB_ACh_) and jRGECO1a were co-excited at 990 nm using an Ultra II femtosecond Ti:Sapphire laser (Coherent) (**Suppl Fig. 4**). Emitted light was filtered (GRAB: 525/50 nm; jRGECO1a: 617/73 nm) and directed to cooled GaAsP detectors (Hamamatsu, H7422P-40). For imaging of vasodilation, Alexa Fluor 680 was conjugated to amino-dextran (MW = 2 MDa, Finabio AD2000x100) (28) and injected IV (50-100 μl of 5 % (w/v) solution in saline). Alexa Fluor 680 was excited using the same 990 nm beam and imaged using a multialkali PMT (R3896, Hamamatsu).

We used a 4x objective (Olympus XLFluor4x/340, NA=0.28) to obtain low-resolution images of the exposure. The Olympus 20x (XLUMPlanFLNXW, NA=1.0) water-immersion objective was used for high-resolution imaging. Measurements were performed in a frame-scan mode at 5-10 Hz.

### Immunohistochemistry

Mice were sacrificed and transcardially perfused with heparinized phosphate-buffered saline (PBS) followed by 4% paraformaldehyde (PFA). The brain was removed immediately after perfusion and post-fixed in 4% PFA overnight at 4°C. The brain was then washed 3 times with PBS and stored at 4°C in a 30% sucrose solution for at least 3 days. Brain tissue was cut into 40 µm coronal or sagittal sections using a cryostat (Leica CM1950) and stored in tris-buffered saline (TBS) containing 0.01% sodium azide at 4°C. Tissue sections were processed for immunofluorescence staining using free-floating staining protocols as previously described (29). In short, antigen retrieval was performed by incubating tissue sections in a 1 N hydrochloric acid solution for 10 min followed by three washes with TBS. Sections were blocked and permeabilized in TBS containing 5% donkey serum and 0.5% triton X-100 for one hour. The primary antibody solution was applied to tissue sections overnight at 4°C and consisted of chicken anti-GFP (1:1000; Abcam, ab13970) diluted in TBS buffer. Tissue sections were then incubated for 2 hours in the secondary antibody solution consisting of Alexa Fluor® 488 donkey anti-chicken (1:250; Jackson ImmunoResearch, AB_2340375), diluted in TBS containing 5% donkey serum. Cell nuclei were stained with 4’,6’-diamidino-2-phenylindole dihydrochloride (DAPI; 2 ng/mL; Molecular Probes) for 15 minutes. Tissue sections were mounted onto slides using ProLong Gold anti-fade reagent (InVitrogen, Cat# P36934) and imaged in the epifluorescence mode on an IX83 Olympus Microscope.

### Data analysis

Mesoscopic data pre-processing and hemodynamic correction of the fluorescence channels were performed as described in (18). Briefly, the modified Beer-Lambert law was used to calculate changes in HbO and HbR concentrations from 525 nm and 625 nm reflectance. Estimated Δ[HbO] and Δ[HbR] were then used to correct the green (GRAB_NE_/GRAB_ACh3.0_) and red (jRGECO1a) channels (19, 27). Mesoscopic images were smoothed in space using a Gaussian kernel with FWHM of 2 pixels.

To calculate the eye pupil diameter, an elliptical region was drawn defining the eye and an intensity threshold was defined as the average of the intensity of the pupil and the surrounding iris. This pupil size was calculated as the number of pixels below this threshold and adjusted to represent pupil diameter in reference to the visible diameter of the eye. To quantify whisking, a rectangular region was drawn over the whisker pad and a motion energy algorithm was used to estimate whisking behavior (30).

### Global IRF model

Prior to IRF estimation, HbT signals were low pass filtered below 0.5 Hz using a Butterworth filter (MATLAB) and Ca^2+^ signals were corrected for the hemodynamic artifact (as described above). Next, HbT and Ca^2+^ signals were normalized, pixel-wise, to their standard deviation in time to remove bias towards pixels with larger variance. With a global, stationary IRF model, HbT is expressed as follows:

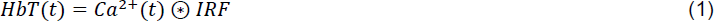

where IRF is invariant with respect to location within the exposure or time. To estimate the IRF, we deconvolved the HbT signal with the Ca^2+^ signal using the matrix formulation below where T is the number of time points and N is the number of pixels:

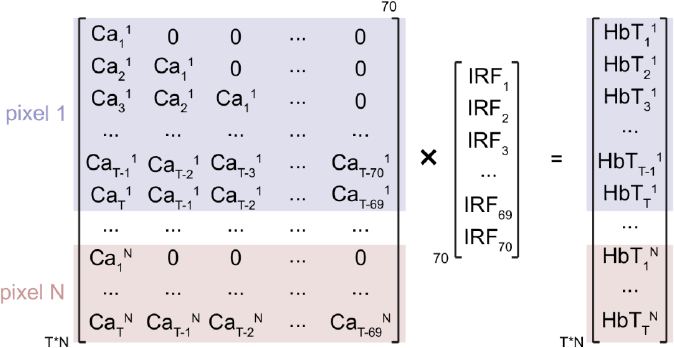

We assumed the following functional form of IRF adopted from (31):

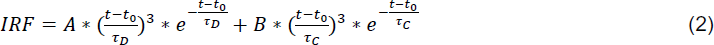

where t_0_ ≥ 0, A and B are scalar coefficients, and *t_D_* and *t_C_* are dilation and constriction time constants, respectively. We simultaneously fit for 5 parameters (t_0_, A, B, *t_D_*, *t_C_*) in MATLAB (R2022a) using the *fminsearch* function to obtain a 7-s (70 time points given 10 Hz acquisition) IRF.

To evaluate the accuracy of this model across cortical space, we convolved, pixel-by-pixel, the estimated global IRF with the Ca^2+^ signal to predict HbT. We then computed, pixel-by-pixel, correlation between the predicted and experimentally obtained HbT using the entire duration of data run (10 min).

To explore dependence of the model accuracy on neuromodulation dynamics, we used a 15-s sliding window to compute (a) correlation between the estimated and experimentally obtained HbT, (b) mean eye pupil diameter, and (c) mean NE signal (averaged across space and time). The latter was motivated by observing that NE was largely uniform across the cortex. Then, for each 15-s time window, we plotted the correlation between the estimated and experimentally obtained HbT as a function of the mean eye pupil diameter or NE in a scatter plot (**Suppl Mov.2**).

### Linear regression model

In our Linear regression model, HbT is expressed as follows:

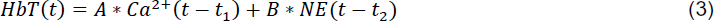

where *t_1_* and *t_2_* are global parameters representing, respectively, the delay between each of the neuronal variables (Ca^2+^ and NE) and HbT, and A and B are scalar weights that can vary across space (from pixel to pixel).

HbT signals were low pass filtered below 0.5 Hz using a Butterworth filter. Ca^2+^ and NE signals were corrected for the hemodynamic artifact (as described above). Next, HbT, Ca^2+^ and NE signals were normalized, pixel-wise, to their standard deviation in time. To find t_1_ and t_2_, we calculated Ca^2+^/HbT and NE/HbT lag cross-correlation functions. This analysis revealed that HbT typically lagged Ca^2+^ and NE by ∼0.9 s and ∼0.1 s, respectively. Therefore, we set t_1_=0.9 s and t_2_=0.1 s.

Next, to calculate the linear regression coefficients, A and B, we performed linear regression, pixel-by-pixel, using the backslash operator in MATLAB. As with the global IRF model, we quantified the accuracy of the Linear regression model as the correlation between the predicted and experimentally obtained HbT.

### Double IRF model

In our Double IRF model, HbT is expressed as follows:

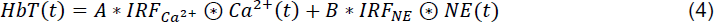

where IRF_Ca2+_ and IRF_NE_ are two stationary IRFs with respect to Ca^2+^ and NE, and A and B are scalar weights that can vary pixel to pixel.

As with the Linear regression model, we low pass filtered the HbT signal below 0.5 Hz and normalized HbT, Ca^2+^, and NE signals, pixel-wise, to their standard deviation in time. Because HbT typically lagged NE by only ∼0.1 s, we chose an IRF kernel ranging from -3 to 7 s. IRF_Ca2+_ and IRF_NE_ were obtained by blind deconvolution using the following design matrix:

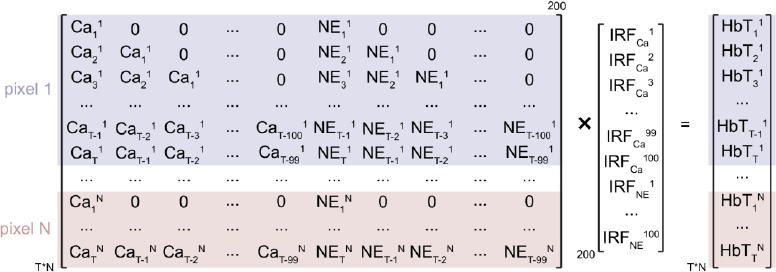

The scalar coefficients, A and B, were estimated, pixel-by-pixel, by regressing the resulting IRF_Ca2+_⊗Ca^2+^ and IRF_NE_⊗NE from HbT using the backslash operator in MATLAB.

### FC Analysis

First, we low pass filtered the HbT signal below 0.5 Hz and spatially downsampled the signals by a factor of 4 using a custom MATLAB function. Then, we calculated seed correlation matrices for HbT and Ca^2+^ using a 30-s sliding window (**Suppl Mov.3**). To compare the HbT and Ca^2+^ FC patterns across time, we shifted the Ca^2+^ time series to precede that of HbT by 0.9 s to account for the neurovascular delay and calculated the correlation between the HbT and Ca^2+^ seed correlation matrices.

To relate the similarity between the Ca^2+^ and HbT connectivity patterns to NE release, we computed NE time-course by averaging the NE signal across all pixels within the exposure at every frame (i.e., time point). We then calculated the correlation between the NE time-course and the time-varying correlation between the Ca^2+^ and HbT seed correlation matrices.

For cortical region-based analysis, we first parcellated the cortical surface based on the mouse cortical Allen atlas (32) and calculated the average Ca^2+^ and HbT time-course within each parcel. We then calculated the seed correlation matrix using a 30-s sliding window. To examine the dependence of inter-regional connectivity of either Ca^2+^ or HbT on NE, for each inter-regional pair, we plotted the mean connectivity with the 30-s time window against mean NE within the same time window as a scatter plot and computed the slope of the scatter plot.

### Spectral Analysis

We performed all spectral analysis using the *mtspectrumc* function of the Chronux toolbox (33) to calculate the Ca^2+^, ACh, NE, and HbT frequency spectra using 9 tapers and a time bandwidth product of 5. All calculated spectra were normalized to their sum. We used the *coherencyc* function (Chronux) to calculate the coherence between Ca^2+^/HbT, ACh/Ca^2+^, and NE/HbT using 9 tapers and a time bandwidth product of 5.

### Statistics

All data analysis and tests for statistical significance were calculated in MATLAB. Results were averaged across runs for each mouse and then averaged across mice. Errors are shown as the standard error across subjects. Differences between groups were tested using a Welch’s t-test using Satterthwaite’s approximation for the effective degrees of freedom (MATLAB *ttest2* function). Number of mice is given in figure legends and **Suppl. Table 1**.

## Results

### Neurovascular coupling depends on neuromodulation and behavior

We used mesoscopic imaging to simultaneously measure Ca^2+^, hemodynamics (HbO/Hb/ HbT) and release of a neuromodulatory transmitter (NE or ACh) over the dorsal cortical surface in fully awake mice (**Fig. 1A-B** and **Suppl Movie 1**; see **Methods**). First, we focused on the relationship between Ca^2+^ as a measure of local neuronal activity and HbT – a measure reflecting vasodilation/constriction (under the assumption of constant hematocrit). Applying a classical hemodynamic response model, where a global (stationary) IRF (see **Methods**), is convolved with the neuronal variable (Ca^2+^) to return the hemodynamic variable (HbT), produced poor results (**Fig. 1C**). Estimating IRF for the medial part of the somatosensory regions of interest (ROI), where the model’s performance was better compared to other regions, further improved the fit for that ROI but at the cost of reduced accuracy elsewhere (**Fig. 1D**).

A close inspection of the Ca^2+^ and HbT time-courses derived from different ROIs revealed instances of divergent HbT despite coherent Ca^2+^ activity (**Fig. 1E**) arguing against a space-invariant convolution kernel. Critically, the accuracy of the convolution model decreased with arousal based on the eye pupil diameter (see **Methods**, **Fig. 1F**).

The eye pupil diameter, whisking and body motion were highly correlated among themselves (**Suppl Fig. 1A**) and with the level of NE (**Suppl Fig. 1B**). Therefore, the model accuracy negatively scaled with NE (**Fig. 1F** and **Suppl Movie 2**). In contrast, there was only weak dependence on the level of ACh (**Suppl Fig. 1C-D**). Because mice in this study expressed either GRAB_NE_ or GRAB_ACh_, the analysis with respect to NE and ACh was performed in a different group of subjects. ACh differed from NE in that it had high similarity with low pass filtered Ca^2+^ (< 0.5 Hz) (**Suppl Fig. 1E-H**).

Given vasoactive properties of NE and the inverse dependence of the stationary IRF model accuracy on the NE level, these results put forward NE as a potential missing link affecting the relationship between the observed spontaneous Ca^2+^ signals and hemodynamics, possibly with some degree of spatial specificity.

### Accounting for NE dramatically improves hemodynamic estimation

Mesoscopic Ca^2+^ signals imaged from the cortical surface are dominated by Ca^2+^ electrogenesis in the apical dendrites of cortical pyramidal cells (PCs) (34). These signals are slower than spike-induced Ca^2+^ transients in neuronal cell bodies due to (1) long duration of dendritic Ca^2+^ spikes and (2) only partial synchronization of activity in a population of PCs. Therefore, although neuronal electrical activity is much faster than hemodynamics, mesoscopic Ca^2+^ signals and hemodynamics have significant overlap in their power spectrum (**Suppl Fig. 2A-B**). We exploited this feature of mesoscale Ca^2+^ data as well as the slow nature of the NE signal to devise our Linear regression model where HbT time-course was predicted, pixel-by-pixel, by a weighted sum of Ca^2+^ and NE time-courses (**Fig. 2**). Assuming the causal neurovascular relationship, the hemodynamic response follows neuronal activity. Therefore, we allowed for a fixed delay between the neuronal variables and HbT estimated by lag cross correlation of HbT with Ca^2+^ or NE (see **Methods**; **Suppl Fig. 2C**). We further reasoned that the contribution of Ca^2+^ and NE to HbT fluctuations can vary in space. Therefore, we introduced regression weights A (for Ca^2+^) and B (for NE) for each pixel; each one can be viewed as a map (**Fig. 2A-B**; see **Methods**). With this model, A weights were consistently positive indicating an increase in HbT, i.e., vasodilation. Conversely, B weights were consistently negative indicating a decrease in HbT, i.e., vasoconstriction (**Fig. 2B**).

**Figure 2:**
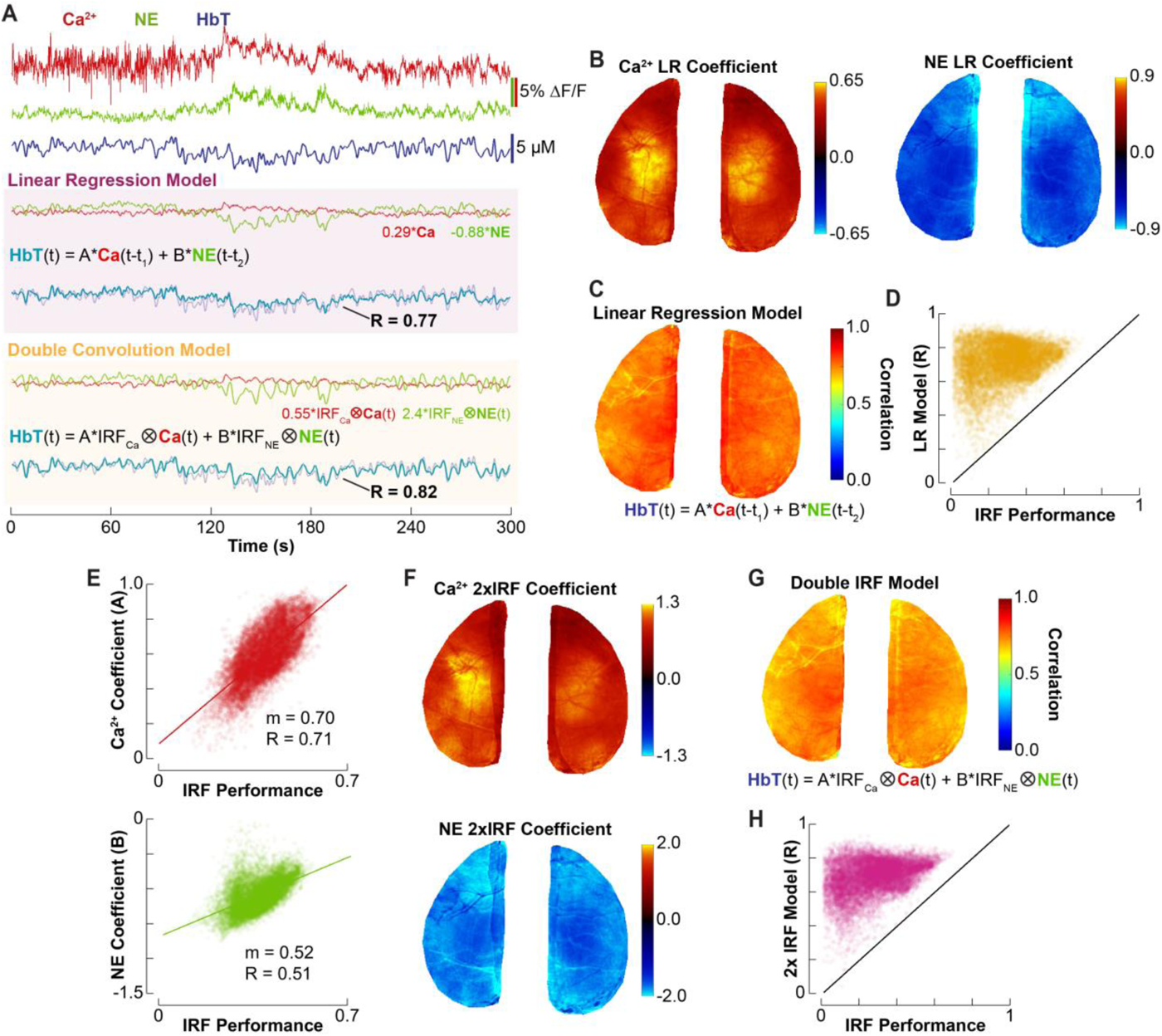
Spontaneous hemodynamic fluctuations depend on both Ca^2+^ and NE dynamics. **A)** Top: example Ca^2+^, NE, and HbT time-courses derived from MOs cortical region. Middle: Linear regression model; time-courses of Ca^2+^ and NE multiplied by the respective weights and the comparison between experimental and predicted HbT (light blue and gray respectively, R=0.77). Bottom: Double IRF model; Ca^2+^ and NE convolution products multiplied by the respective weights and the comparison between the experimental and predicted HbT (light blue and gray respectively, R=0.82. **B)** Maps of the weights, A (left) and B (right), in the Linear regression model (averaged over n=7 mice). **C)** Stationary IRF accuracy map quantified as the correlation between experimental and predicted HbT (averaged over n=7 mice). **D)** Pixel-by-pixel accuracy of the Linear regression model plotted against that of the stationary IRF model along with the line of equal performance (n=1 mouse). **E)** Pixel-by-pixel weights for Ca^2+^ (A) and NE (B) plotted against the pixel-by-pixel stationary IRF performance (n=1 mouse). **F)** Maps of the weights, A (top) and B (bottom), in the Double IRF model (n=7 mice). **G)** Same as (C) for the Double IRF model (n=7 mice). **H)** Same as (D) but for the Double IRF model (n=1 mouse).

Compared to the global IRF model, the Linear regression model dramatically improved the accuracy increasing the mean correlation (for all pixels) between the predicted and experimentally obtained HbT from 0.22±0.05 to 0.78±0.01 (n=7 subjects, 11 hemispheres; **Fig. 2C-D** and **Suppl Fig. 3**). Notably, the regression weights for the Ca^2+^ signal were high in cortical areas where performance of the global IRF was fair and low in those regions where the global IRF model performed poorly (**Fig. 2E**). The regression weights for the NE signal had a weaker spatial relationship but were typically more negative in regions where performance of the global IRF was poor (**Fig. 2D**, **Suppl. Fig. 2E**). Similar but slightly weaker results were obtained when HbT was substituted by HbO and Hb (HbO: 0.68±0.04; Hb: 0.53±0.05; **Suppl Fig. 2F**).

Conceptually, these results suggest two parallel (non-interacting) forces governing HbT behavior that are captured by Ca^2+^ (dilation) and NE (constriction) signals – the NE signal has a negative correlation with hemodynamic fluctuations as well as vessel diameter fluctuations observed in 2-photon experiments (**Suppl Fig. 4**). The influence of each of these forces on HbT varies across space. Therefore, we decided to explore whether the accuracy of the IRF model can be improved by incorporating two stationary IRFs – IRF_Ca2+_ and IRF_NE_ – along with space-varying weights A and B reflecting that contribution of Ca^2+^ and NE to HbT can vary in space. To devise this Double IRF model, we used blind deconvolution to estimate IRF_Ca2+_ and IRF_NE_ from the data (see **Methods**). The obtained IRF_Ca2+_ was consistently positive (i.e., vasodilation), while IRF_NE_ – consistently negative (i.e., vasoconstriction; **Suppl Fig. 2D**).

Similar to Linear regression model, Double IRF model dramatically improved the prediction accuracy over the original global IRF model from 0.22±0.05 to 0.74±0.01 (n=7 subjects, 11 hemispheres; **Fig. 2A,G,H** and **Suppl Fig. 3**). Similar to Linear regression model, weights for the Ca^2+^ signal scaled with the performance level of the global IRF model across space whereas weights for the NE signal in space weakly depended on the global IRF performance (**Fig. 2F** and **Suppl Fig. 2E**). Similar but slightly weaker results were obtained when HbT was substituted by HbO and Hb (HbO: 0.71±0.02; Hb: 0.65±0.03; **Suppl Fig. 2G**).

These results demonstrate that spontaneous hemodynamic fluctuations can be captured by a simple linear model with effective parameters describing the (dynamics of) dilatory and constrictive forces while also accounting for their varying influence across space.

### Hemodynamic FC is a poor predictor of neuronal FC during arousal

The spatial specificity of the NE effect on hemodynamics implies that, for some cortical regions, reduction in the hemodynamic coherence during arousal can occur despite coherent behavior of the underlying neuronal circuits. Indeed, close inspection of our data revealed instances of divergent hemodynamic activity for pairs of cortical regions during high arousal (high NE and large eye pupil diameter) in the presence of coherent Ca^2+^ activity (e.g., between the visual and motor cortices (VISp and MO, **Fig. 3A**).

**Figure 3:**
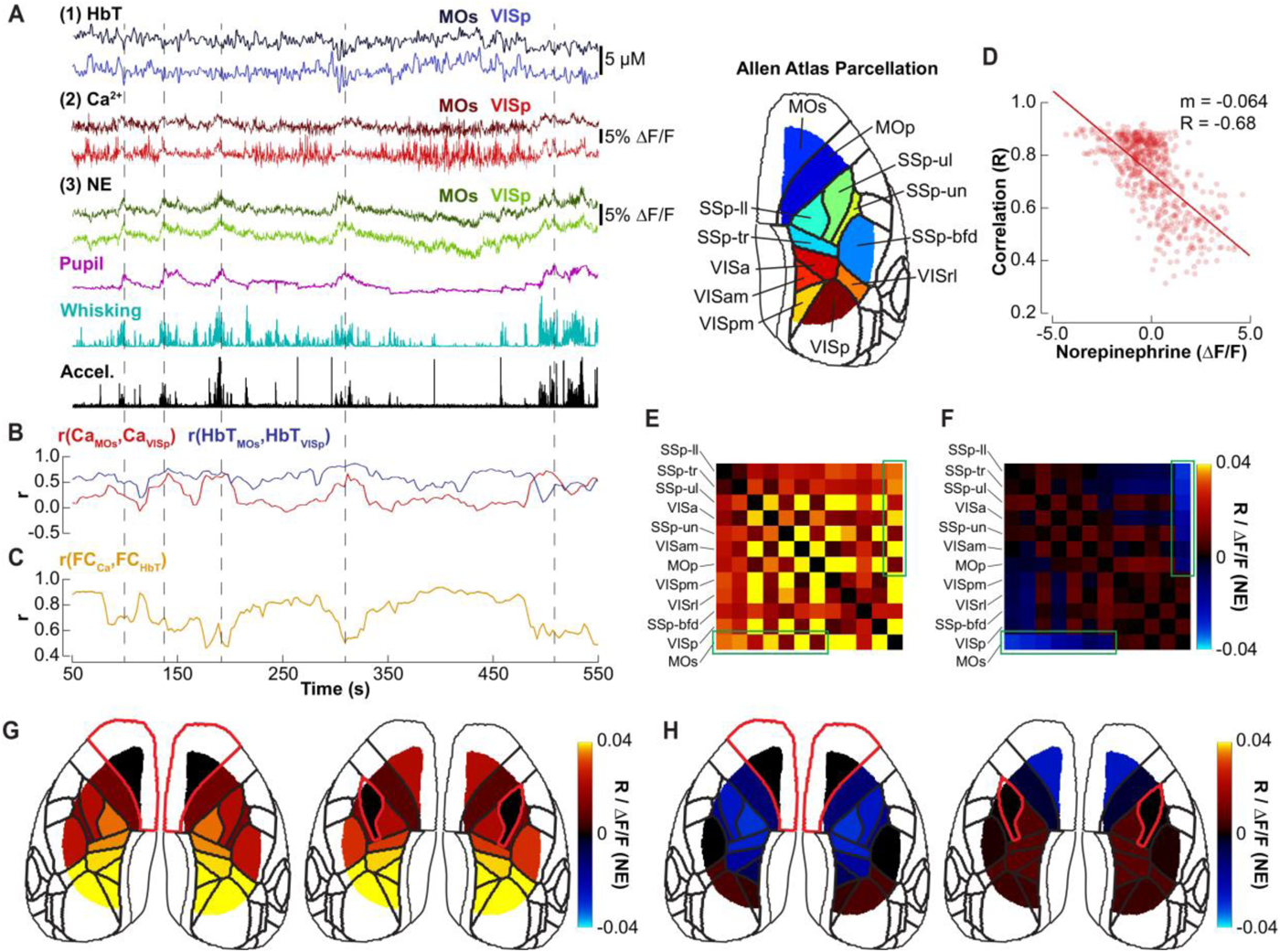
FC patterns between HbT and Ca^2+^ diverge during high arousal. **A)** Example Ca^2+^, NE, and HbT time-courses derived from two cortical regions, MOs and VISp, along with behavior readouts. **B)** Time-varying inter-regional correlation between MOs and VISp for Ca^2+^ (red) and HbT (blue). **C)** Time-varying correlation between FC patterns of Ca^2+^ and HbT. **D)** Same data as in (C) shown as a function of NE. Each point on the scatter plot corresponds to a correlation coefficient for a 30-s sliding window plotted against the NE within the same time window. The slope shows a strong negative relationship. **E)** For each cortical region pair, we computed the slope of a scatter plot of the Ca^2+^ connectivity vs. NE; this slope is color-coded in the matrix (average of n=7 mice). **F)** Same as (E) for HbT (average of n=7 mice). **G)** One row of (E) – MOs (left) and SSp-II (right) plotted as a cortical map. **H)** One row of (F) – MOs (left) and SSp-II (right) plotted as a cortical map.

To quantify temporal dynamics of this phenomenon with respect to NE, we first calculated FC as the pixel-wise seed-correlation matrices using a 30-second sliding window for each Ca^2+^ and HbT time series (FC_Ca2+_ and FC_HbT_, **Fig. 3B** and **Suppl Movie 3**). Then, we performed correlation between the resulting matrices (r(FC_Ca2+_, FC_HbT_); see **Methods**). As expected, this analysis revealed departures between FC_Ca2+_ and FC_HbT_ during high alertness (high NE and large eye pupil diameter) with an overall negative correlation between r(FC_Ca2+_, FC_HbT_) and NE (**Fig. 3C-D** and **Suppl Movie 3**). A similar relationship was found when only high frequency (0.5-5 Hz) Ca^2+^ was considered (**Suppl Fig. 5**).

Next, we examined the regional specificity of the relationship between FC_Ca2+_ and FC_HbT_ with respect to NE. To this end, we used the Allen atlas (32) to define 12 primary cortical regions within the exposure. For every pair of cortical regions, we used a 30-second sliding window to calculate correlation between Ca^2+^ time-courses extracted from these regions and plotted the correlation value as a function of the mean NE within the same time window in a scatter plot. Then, we computed the slope of the scatter plot. The value of this slope for every pair of cortical regions is shown in **Figure 3E**. The same procedure was repeated for HbT (**Fig. 3F**). Visual comparison of **Figures 3E** and **F** reveals anticorrelation between some pairs of cortical regions upon an increase in NE (blue colors in **Fig. 3F**). In contrast, correlation between Ca^2+^ signals increased or remained the same (red and yellow colors in **Fig. 3E**). In particular, we observed a robust hemodynamic anticorrelation between anterior cortical areas (MOs) and somatosensory areas (outside barrel cortex) in the presence of high NE, which occurred despite an increase in Ca^2+^ correlation (green rectangles in **Fig. 3E-F**). These results are consistent with the results in **Figure 2** showing high NE and low Ca^2+^ weights for HbT fit in the anterior regions (i.e., dominant negative IRF_NE_) and the opposite – high Ca^2+^ and low NE weights for HbT fit in the somatosensory areas (i.e., dominant positive IRF_Ca2+_). The same finding is illustrated in the seed connectivity maps in **Figure 3G-H**.

Thus, vasoconstriction upon an increase in arousal and NE resulted in flipping of neurovascular coupling (NVC) with respect to neuronal Ca^2+^ from positive to negative in the anterior but not somatosensory cortex and anticorrelation of the hemodynamic signals between these cortical regions. On the other hand, the correlation of Ca^2+^ activity between cortical regions in general increased upon an increase in the arousal/NE.

## Discussion

Taken together, our results show that hemodynamic fluctuations accompanying spontaneous neuronal activity strongly depended on NE neuromodulation. While NVC with respect to neuronal Ca^2+^ was a dynamic and seemingly complex phenomenon, modeling the hemodynamic signal as a weighted sum of either (a) Ca^2+^ and NE time-courses with the Linear regression model, or (b) stationary IRF_Ca2+_ and IRF_NE_ convolved with the respective Ca^2+^ and NE time-courses with the Double IRF model, produced a high accuracy fit to the data (0.76±0.02 and 0.72±0.02 accuracy, respectively). While this behavior was common for HbT, HbO and HbR, we discuss our results with respect to HbT that is related to the arteriolar diameter, the physiological parameter directly controlled by vasoactive messenger molecules. Prior work on vascular responses evoked by sensory and OG stimulation by us and others demonstrated that dilatory and constrictive forces co-exist and often result in a biphasic behavior of cortical arterioles (31, 35, 36). In the current study, these forces were captured by the Ca^2+^ (dilation) and NE (constriction) coefficients, in both the Linear regression and Double IRF model. The high accuracy of HbT prediction with these models suggests that feed-forward, neurogenic regulation of CBF, which governs stimulus-induced hemodynamic responses (35, 37–39), also applies to spontaneous (resting-state) neuronal activity. Therefore, analysis of resting-state (as well as single-trial) hemodynamic data should consider fluctuations in the arousal affecting NVC and IRF, and a failure to account for these effects may result in erroneous interpretations of hemodynamic coherence, or lack thereof, in terms of the underlying neuronal activity. Our results also imply that non-neurogenic forces such as intrinsic vasomotion do not significantly contribute to resting state hemodynamics.

Like other vasoactive neuromodulatory neurotransmitters, NE affects NVC in two ways: (a) directly, by acting through adrenergic receptors on the vascular smooth muscle cells, and (b) indirectly, by modulating the local neuronal circuit activity and the associated release of vasoactive messengers (4). Earlier studies that employed electrical, OG or pharmacological stimulation of LC produced controversial results regarding dilation/constriction effects in cerebral cortex (40, 41). This is in contrast with tissues outside the brain where activation of vascular adrenergic receptors consistently produces vasoconstriction (42). In this respect, it is worth noting that stimulation of LC can increase neuronal activity (43) and that release of vasodilators from activated local cortical neurons can mask vasoconstriction due to activation of vascular adrenergic receptors. In contrast to these perturbation approaches, we used a fluorescent biosensor GRAB_NE_ (14) to observe the release of NE in the context of naturally occurring brain activity. Under these conditions, NE was clearly correlated with a decrease in HbT in mesoscopic imaging and vasoconstriction in 2-photon arteriolar diameter measurements (**Suppl Fig. 4**).

Meso-scale NE signals had low spatial granularity, which is in agreement with previous reports on the diffuse nature of NE projections in cerebral cortex (44). The importance of NE for HbT prediction (i.e., pixel-by-pixel weighting), however, varied across the cortical space. The Double IRF model outperformed the canonical convolution model with a global stationary IRF across all pixels, demonstrating a clear dependence of NVC on NE and behavior (the eye pupil diameter, whisking, and body movement). Even in the medial part of the somatosensory cortex, where the contribution of NE to HbT was relatively small, the relationship between Ca^2+^ and HbT was poorly captured by the single IRF model (0.38±0.05 accuracy). The highest contribution of NE to HbT was found in the anterior cortex. These results are in general agreement with previous studies – each one focused on a different brain region – that reported different IRFs (39, 45).

Because of the low spatial granularity of NE signals, the differences in NE contribution to HbT across cortical regions likely arose due to postsynaptic mechanisms including the type and density of (neuronal, glial, and vascular) adrenergic receptors as well as the density of target cortical neurons, e.g., inhibitory interneurons expressing a vasoconstrictor Neuropeptide Y (46).

Given the prevalence of α-adrenergic receptors on vascular smooth cells (47, 48), these receptors may mediate the direct vascular effects of NE.

In contrast to NE, meso-scale ACh signals had high similarity with meso-scale Ca^2+^ signals filtered below ∼0.5Hz, and therefore did not offer an additive value to that of Ca^2+^ signals alone for prediction of HbT. Modeling of light propagation in cortical tissue by us and others has revealed that wide-field fluorescence imaging is mostly sensitive to signals within the top ∼100 μm of cerebral cortex (49), e.g. cortical layer I that houses pyramidal apical dendritic tufts possessing voltage-gated Ca^2+^ channels (34). Because activation of pyramidal neurons elicits vasodilation (35, 50, 51), and ACh is a vasodilator as well (52), their respective vasodilatory effects would be lumped into the Ca^2+^ weighting coefficient in our Linear regression and Double IRF models. In the current study, we made no attempt to untangle these effects. In addition, we did not distinguish between ACh released by BF projections and that released by local cortical ChAT-VIP interneurons (53). Given the presence of local cholinergic axons in cortical layer I (54), high sensitivity of wide-field imaging to the cortical surface (49), and our recent findings of retinotopic ACh responses in the primary visual cortex (55), we believe that both sources contributed to the observed GRAB_ACh3.0_ fluorescence.

In human BOLD fMRI studies, the eye pupil dilation was shown to correlate with deactivation of the default mode network, DMN (56–58). In our data, the anterior cortical region where NE and eye pupil diameter were correlated with a decrease in HbT may belong, at least in part, to the mouse DMN analog (59–62). Although firing rate decreases have been reported in other parts of DMN during (hemodynamic) deactivation (63), the present results suggest that active (signaling-driven) vasoconstriction should also be considered. Similar conclusion was reached in a recent mouse study that examined the effect of tonic chemogenetic activation of LC on frontal cortex part of DMN (8).

The spatiotemporal patterns of FC in resting state BOLD signals have become the principal means to investigate the integrity of large-scale functional networks in the human brain in health and disease (64). Further, the concept of dynamic FC was introduced in humans (65) and validated in animals (66, 67). The main underlying assumption in FC studies is that coherence of hemodynamic signals reflects synchronization of the underlying neuronal dynamics. Consistent with published studies, NE in our data was highly correlated with arousal: an increase in the eye pupil diameter, whisking, and body movement (2). During high arousal, we observed instances of diminished hemodynamic coherence between cortical regions where HbT was differentially impacted by NE (e.g., anterior vs. medial somatosensory cortex) despite coherent behavior of the underlying neuronal Ca^2+^ activity. Thus, the spatial specificity of NE affecting hemodynamics can lead to false interpretation of diminished FC as the underlying neuronal desynchronization during high arousal. Fortunately, high correlation of NE with the eye pupil diameter (68) may allow using the eye pupil diameter as a surrogate for NE in fMRI studies where this measurement is readily attainable (56, 58, 69, 70).

In contrast to a human resting-state fMRI study where subjects are typically instructed not to move, head-fixed mice exhibit intermittent whisking and body movements (71). Further, NE, the eye pupil diameter, whisking, and movement were highly correlated with each other in our data. In other words, higher arousal was invariantly associated with motor behavior. Although we could not dissociate arousal from movement, our data clearly demonstrate that NVC depends on the state of arousal and cortical region.

In our study, we used Ca^2+^ as a measure of the activity of local neuronal circuits. This measure is not available in humans. In humans, BOLD fMRI can be combined with Electroencephalography (EEG) that offers a non-invasive measure of the extracellular electrical potential and has its own advantages, limitations, and measurement theory (72). Therefore, translation of the current results and take-home messages to humans requires tackling the relationship between the mesoscopic Ca^2+^ signals, which have been employed here, and surface electrical potentials that can be used for feedforward calculation of the corresponding EEG (37); this is an ongoing effort in our “collaboratory.”

In the present study, we demonstrate that spontaneous hemodynamic fluctuations can be captured by introducing effective parameters reflecting the underlying dilatory and constrictive forces. In the reality, multiple signaling pathways contribute to each of these forces. Future studies will parse those pathways to offer the cell-type and molecular specificity in the context of brain state and behavior. The most immediate implication of our present study for human resting state fMRI is that diminished FC upon arousal may reflect the spatial specificity of the effect of neuromodulation on IRF rather than desynchronization of local cortical circuits.

## Acknowledgements

We thank Mickey London, Anton Arkhipov, Li-Huei Tsai, and Xue Han for helpful discussions and Tim O’Shea for provision of the epifluorescence microscope. We gratefully acknowledge support from the NIH (BRAIN Initiative U19NS123717, BRAIN Initiative R01NS122742, R01DA050159). Patrick Doran was supported by Ruth L. Kirschstein Predoctoral Fellowships F31NS118949. Patrick Bloniasz was supported by NIH T32NS131178 and NSF Graduate Research Fellowship Program under Grant No. 2234657.

## Author Contributions

**Bradley C. Rauscher, Natalie Fomin-Thunemann**: Investigation, Formal Analysis, Conceptualization, Methodology, Visualization, Writing – Original Draft, Writing - Review & Editing. **Sreekanth Kura**: Formal Analysis, Methodology, Software, Writing - Review & Editing. **Patrick R. Doran**: Methodology, Software, Writing – Original Draft, Writing - Review & Editing. **Pablo D. Perez, Dora Balog, Patrick F. Bloniasz**: Formal Analysis, Conceptualization, Writing - Review & Editing. **Nathan X. Chai, Francesca A. Froio, Andrew Garcia**: Investigation, Formal Analysis, Writing - Review & Editing. **Kate E. Herrema**: Investigation, Visualization, Resources, Writing – Original Draft, Writing – Review & Editing. **Kıvılcım Kılıç, John X. Jiang**: Resources, Writing - Review & Editing. **Scott Knudstrup**: Investigation, Writing - Review & Editing. **Jeffrey Gavornik, David Kleinfeld, Michael Hasselmo, Laura D. Lewis, Emily P. Stephen, David A. Boas, Lei Tian, Gal Mishne, Sava Sakadzic**: Conceptualization, Methodology, Supervision, Writing - Review & Editing. **Martin Thunemann**: Conceptualization, Software, Supervision, Writing – Original Draft, Writing – Review & Editing. **Anna Devor**: Conceptualization, Funding Acquisition, Supervision, Writing – Original Draft, Writing – Review & Editing.

---

**Supplementary Figure 1:**
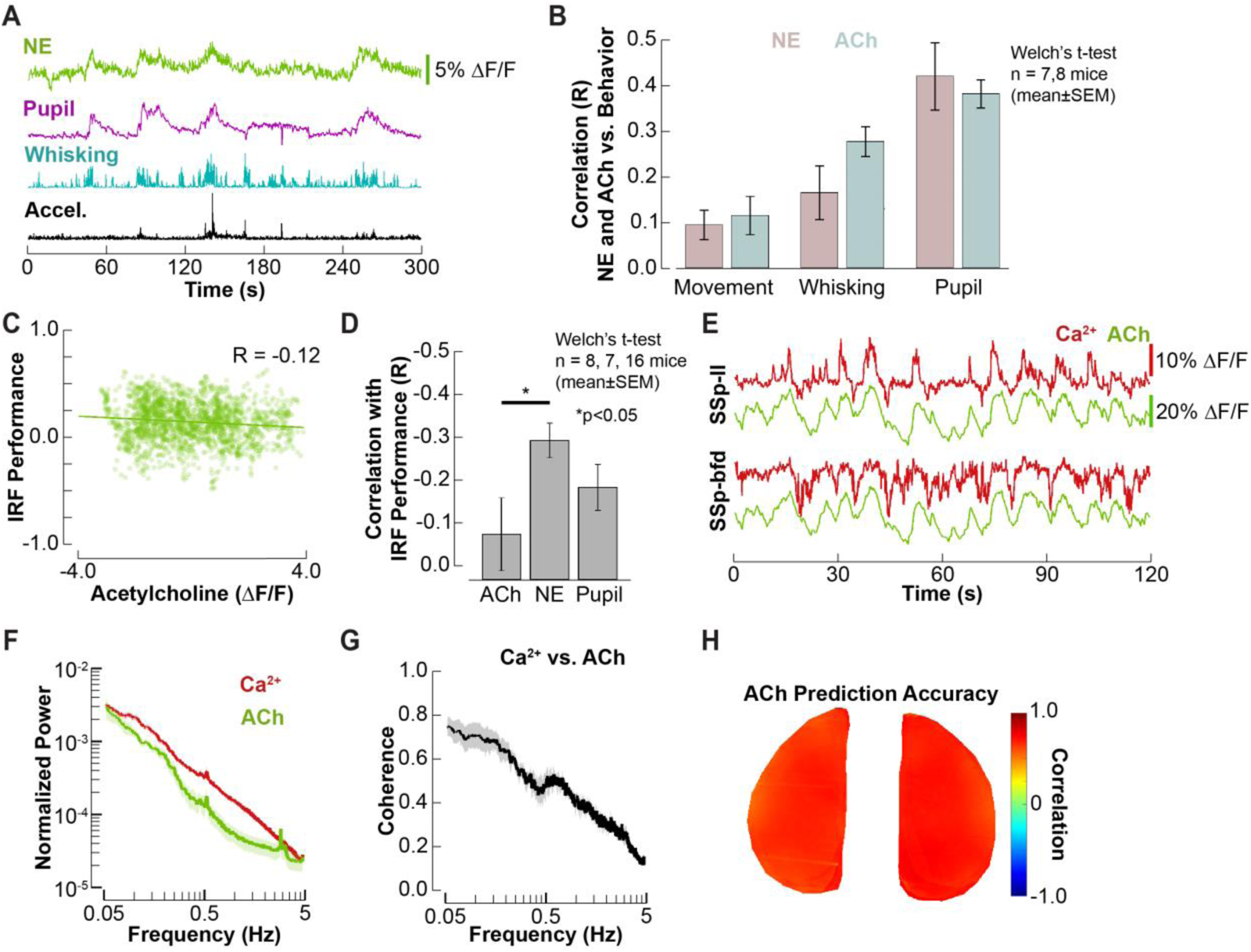
NE and ACh with respect to behavior and high correlation between ACh and low pass filtered Ca^2+^. **A)** Example NE, pupil, whisking, and accelerometer time-courses. **B)** Correlation between NE and ACh with behavioral parameters: movement, whisking, and pupil (averaged over n=7 and 8 mice, respectively; mean±SEM, Welch’s t-test, *p<0.05). **C)** The accuracy of stationary IRF derived from SSp-bfd shows no dependence on ACh (n=1 mouse). **D)** Correlation between the accuracy of stationary IRF derived from SSp-bfd and either ACh, NE, or pupil diameter (averaged over n=8,7,16 mice, respectively; mean±SEM, Welch’s t-test, *p<0.05) **E)** Example Ca^2+^ and ACh time-courses derived from SSp-ll and SSp-bfd. **F)** Normalized power spectra for Ca^2+^ and ACh (averaged over n=8 mice, mean±SEM). **G)** Coherence between Ca^2+^ and ACh (averaged over n=8 mice, mean±SEM). **H)** Prediction accuracy of ACh from low pass filtered Ca^2+^ (<0.5 Hz) quantified as pixel-by-pixel correlation between Ca^2+^ and ACh (averaged over n=8 mice).

**Supplementary Figure 2:**
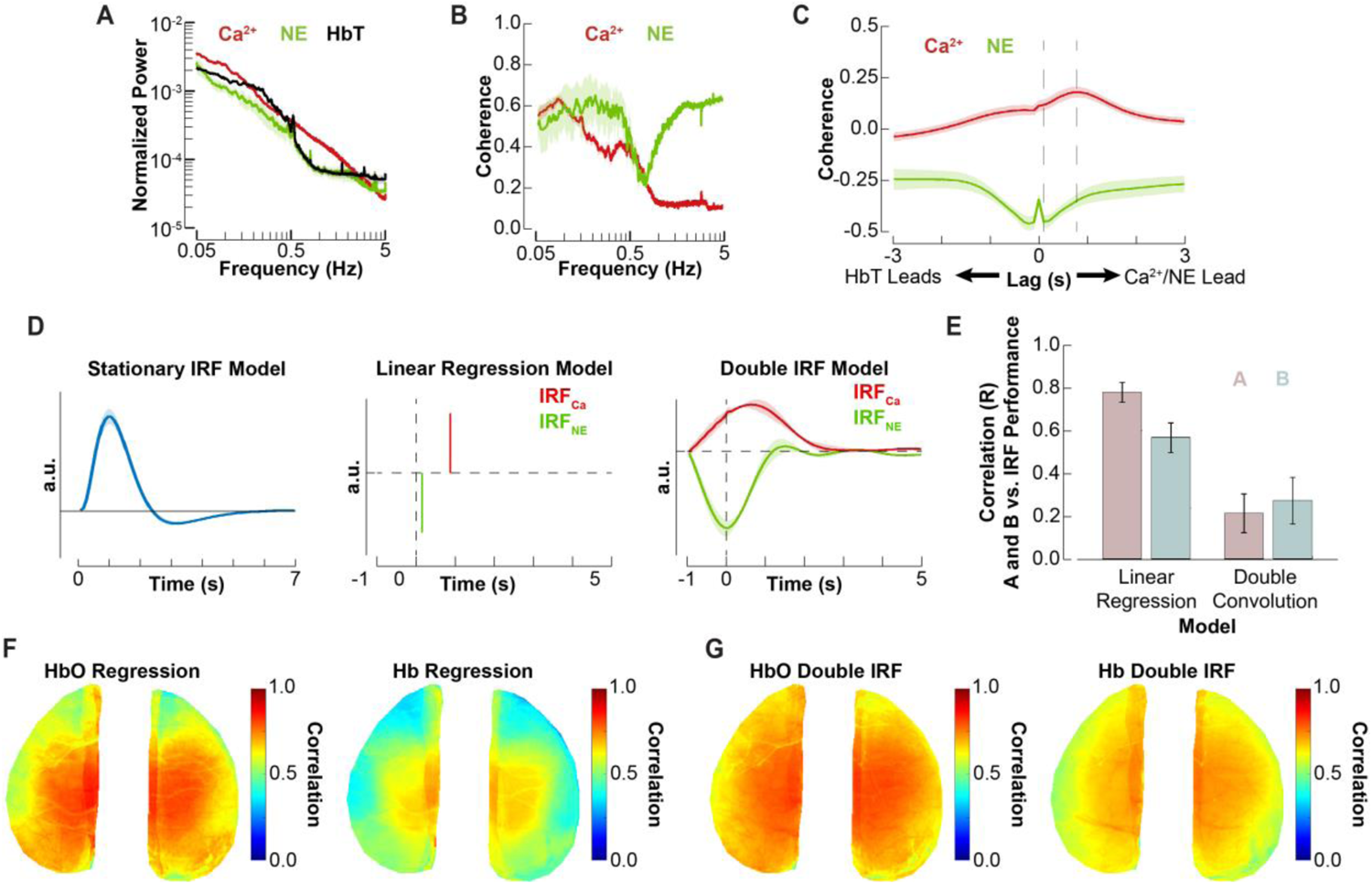
Linear regression and Double IRF prediction models with respect to HbO and HbR. **A)** Normalized power spectra for Ca^2+^, NE, and HbT (averaged over n=7 mice, mean±SEM). **B)** Ca^2+^/HbT and NE/HbT coherence (averaged over n=7 mice, mean±SEM). **C)** Lag cross-correlation between Ca^2+^/HbT (red) and NE/HbT (green) (averaged over n=7 mice, mean±SEM). The sharp peak in the NE/HbT cross-correlation function at t=0 is an artifact of overcorrection with the hemodynamic correction algorithm (see **Methods**). **D)** IRFs for the stationary IRF (n=15 mice), Linear regression, and Double IRF models (n=7 mice) (mean±SEM). **E)** Pixel-wise correlation of A and B weights with stationary IRF accuracy for the Linear regression and Double IRF models (averaged over n=7 mice, mean±SEM). **F)** HbO and HbR prediction accuracy maps for the Linear regression model (averaged over n=7 mice). **G)** Same as (F) but for the Double IRF model (averaged over n=7 mice).

**Supplementary Figure 3:**
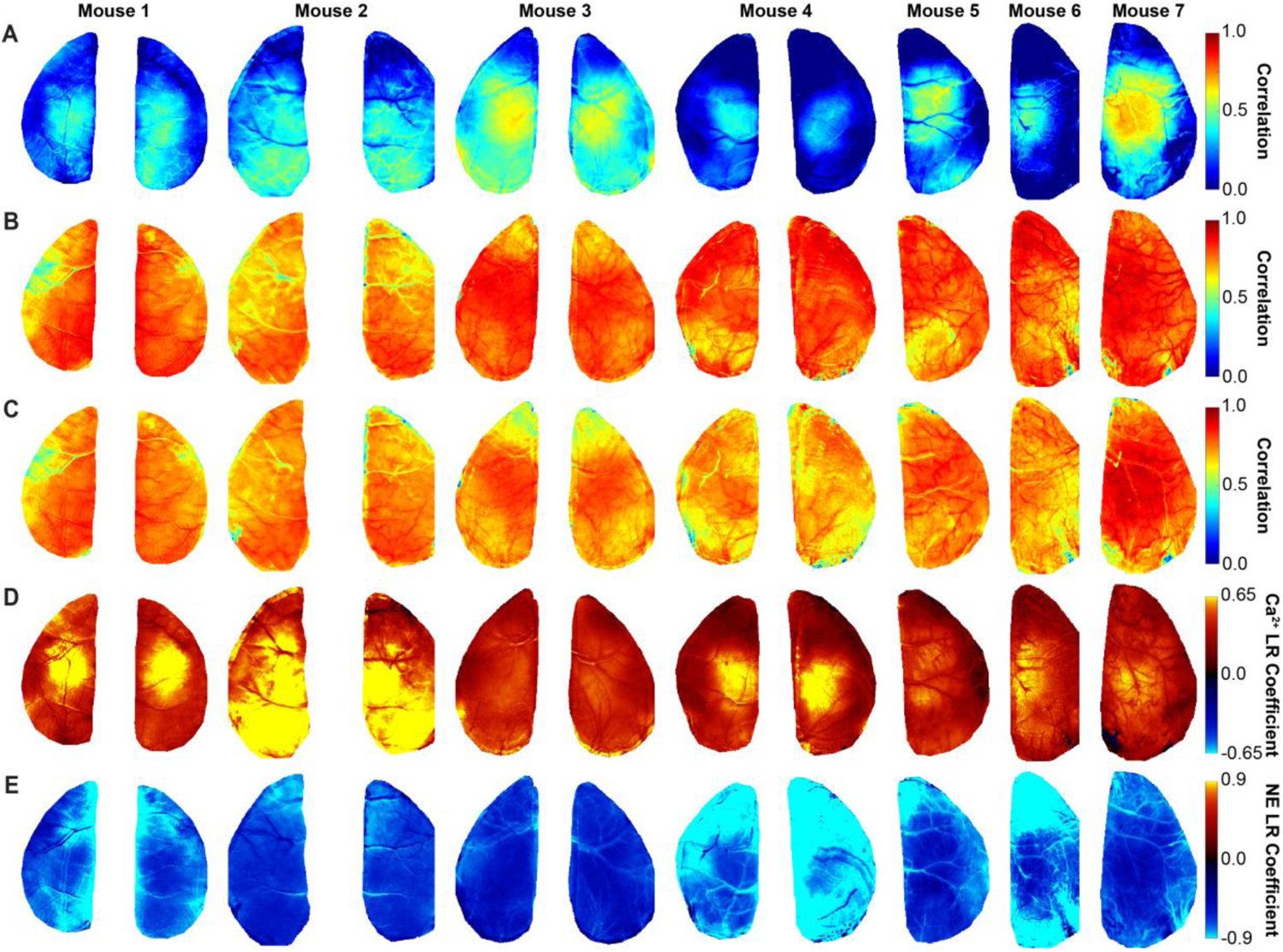
Accuracy of the Linear regression and Double IRF models across mice. **A)** Maps of HbT prediction accuracy of the somatosensory-derived stationary IRF model for each mouse. **B)** Maps of HbT prediction accuracy of the Linear regression model for each mouse. **C)** Maps of HbT prediction accuracy of the Double IRF model for each mouse. **D)** Maps of Ca^2+^ weighting coefficient A for each mouse. **E)** Maps of NE weighting coefficient B for each mouse.

**Supplementary Figure 4:**
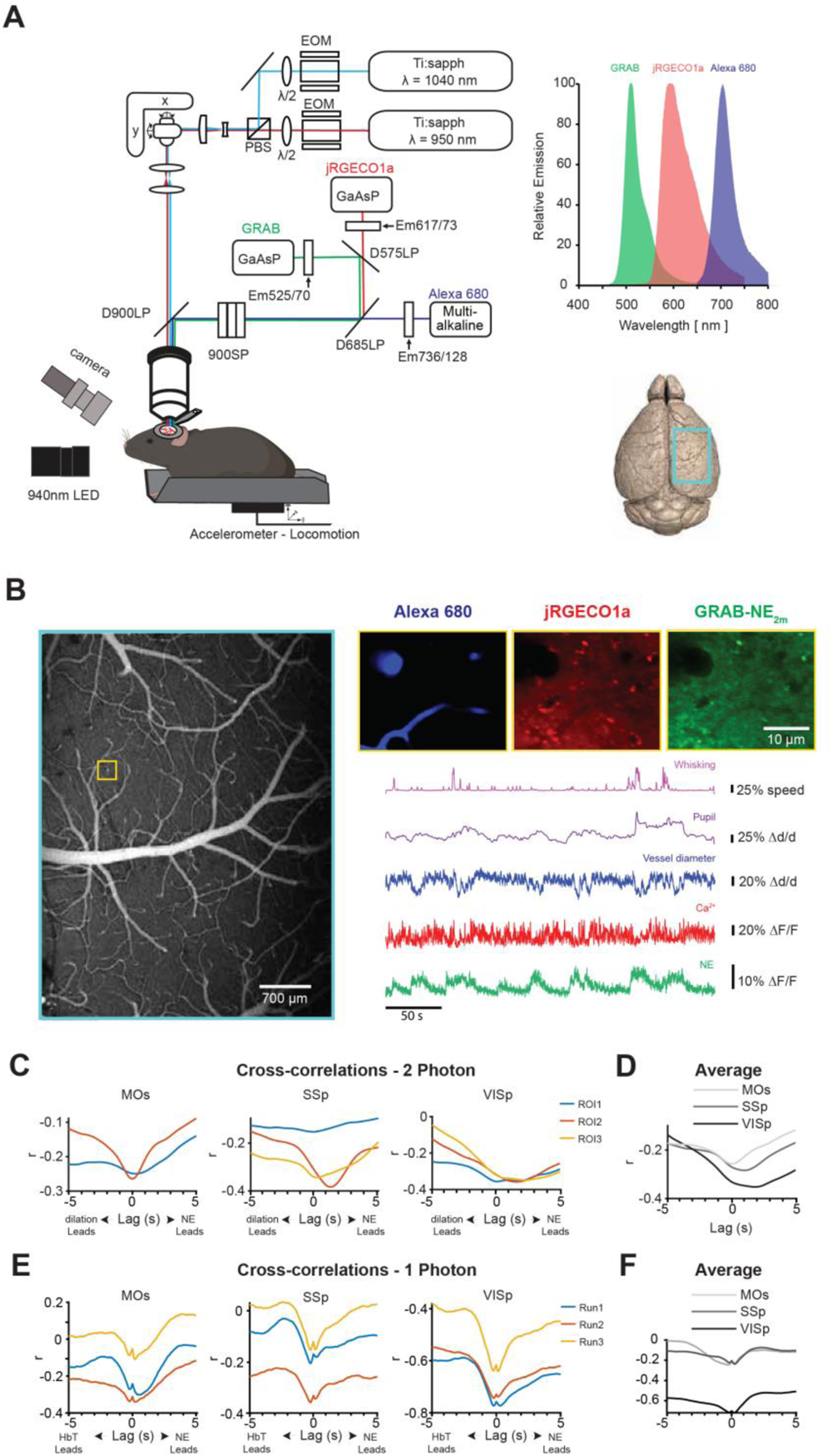
Two-photon imaging of Ca^2+^, NE, and arteriolar diameter. **A**) Imaging setup depicts Ti:sapph laser set to 990 traveling through an electro-optic modulator (EOM) to excite GRAB_NE_, jRGECO1a and Alexa 680; D900LP, D685LP and D575LP—long-pass dichroic mirrors with a cutoff at 900, 685 and 575 nm, respectively; GaAsP and multialkaline— PMT tubes; 900SP—short-pass optical filter with a cutoff at 900 nm; Em525/70, Em617/70, Em736/128—bandpass emission filters. Absorption plot at the top right indicates the emission spectra of our desired fluorophores (spectrum of EGFP-derived GRAB_ACh3.0_ taken from (16); spectrum of mApple-derived jRGECO1a taken from (73); spectrum of Alexa 680 taken from *SpectraViewer* (Thermofisher)). Rendering of the brain surface (taken from (74)) in the bottom right shows the area (turquoise rectangle) imaged in B. **B**) Left: An example average intensity projection of a low magnification image stack across the top 300 µm showing the vasculature labeled with Alexa 680. Yellow rectangle indicates the ROI chosen for imaging of brain activity. Right: A field of view 170 µm below the cortical surface with a diving arteriole labeled with Alexa 680(blue), neurons labeled with jRGECO1a (red), and GRAB_NE_ (green). Time-courses at the bottom show spontaneous activity acquired at 10 Hz. **C**) Cross-correlation of the NE signal and arteriolar diameter for different arteries (ROIs) across three cortical regions: MOs, SSp and VISp. **D**) Average across ROIs for the three cortical regions. **E**) As in (C) for signals derived from mesoscale imaging. **F**) Average across ROIs for the three cortial regions derived from mesoscale imaging.

**Supplementary Figure 5:**
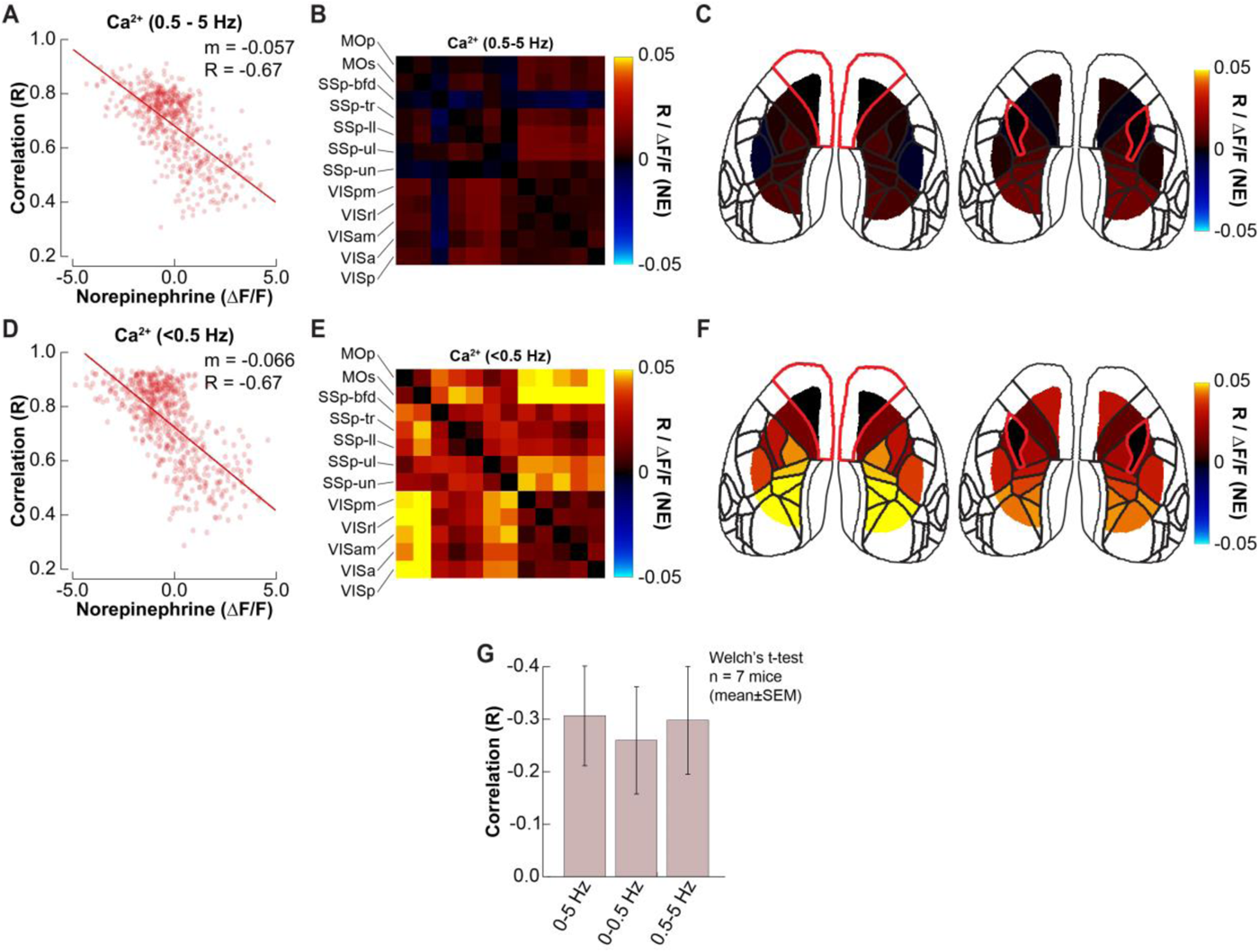
Comparison of FC results for high and low frequency band of Ca^2+^ signal. **A)** Time-varying correlation between FC patterns of high frequency (0.5-5 Hz) Ca^2+^ and HbT plotted as a function of NE. Each point on the scatter plot corresponds to a correlation coefficient for a 30-s sliding window plotted against the NE within the same time window. The slope shows a strong negative relationship. **B)** For each cortical region pair, we computed the slope of a scatter plot of the Ca^2+^ connectivity vs. NE; this slope is color-coded in the matrix (average of n=7 mice). **C)** One row of (B) – MOs (left) and SSpII (right) plotted as a cortical map. **D)** As in (A) for low frequency (0.5-5 Hz) Ca^2+^. **E)** As in (B) for low frequency (0.5-5 Hz) Ca^2+^. **F)** One row of (E) – MOs (left) and SSpII (right) plotted as a cortical map. **G)** Correlation between FC patterns of Ca^2+^ and HbT (r(FC_Ca2+_, FC_HbT_)) with NE for low (0-0.5 Hz) and high (0.5-5 Hz) frequency Ca^2+^ as well as the broadband Ca^2+^ signal (averaged over n=7 mice, mean±SEM).

**Supplementary Figure 6:**
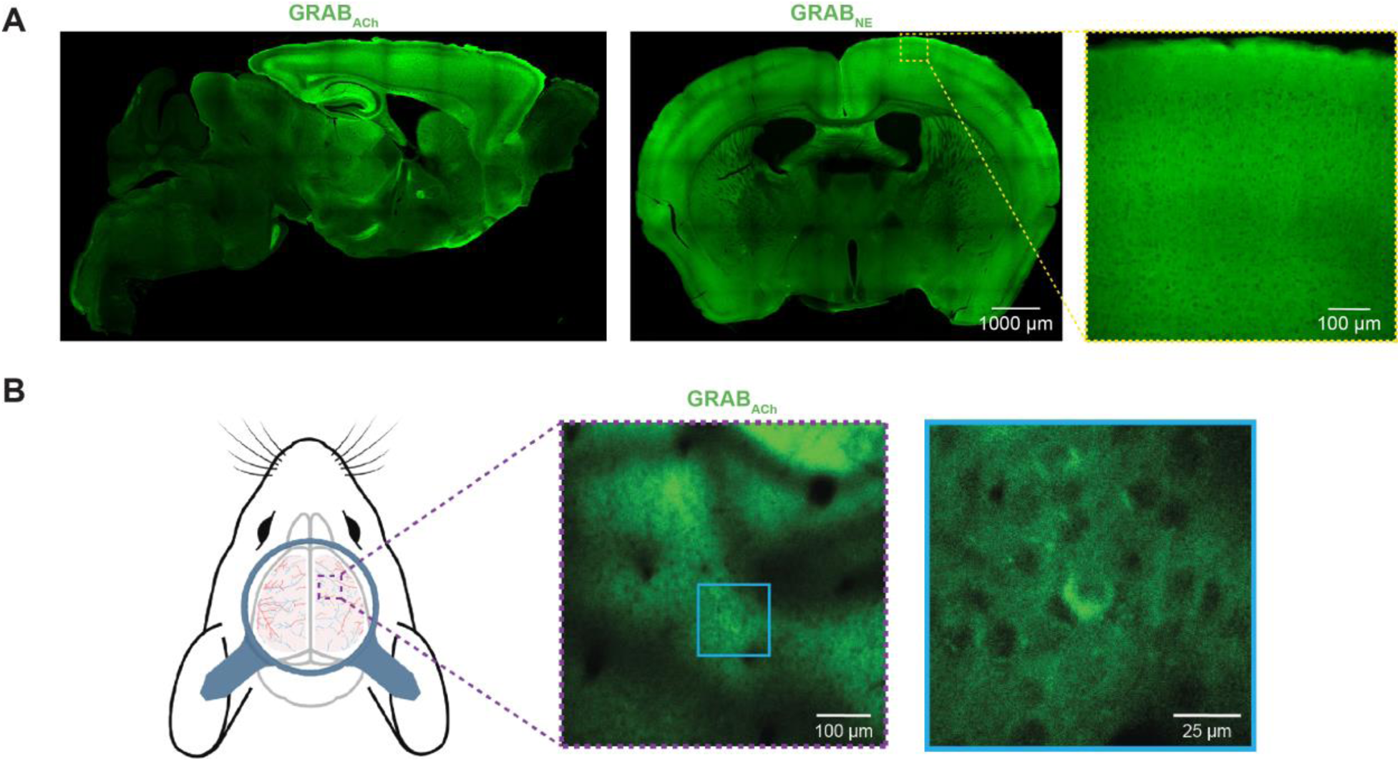
Brain-wide expression of GRAB sensors. **A)** Immunofluorescence staining for GFP of sagittal (left) and coronal (middle, right) mouse brain sections showing GRAB_ACh_ sensor expression at 10x (left) and GRAB_NE_ sensor expression at 10x (middle) and 20x (right) magnification. **B)** Scheme of an awake mouse (BioRender) with head bar and a ROI used to capture 2-photon GRAB_ACh_ fluorescence images (left). Average intensity projection across a 50 µm 3-dimensional stack in layer 2 showing GRAB_ACh_ fluorescence (middle). Enlarged ROI from middle panel (right).

**Supplementary Table 1: Summary of the experimental dataset**

Sensor expressed (GRAB_NE_ or GRAB_ACh_), number of days imaged, and number of minutes imaged for each of the 15 mice in this study.

